# The interplay between Bag-1, Hsp70, and Hsp90 reveals that inhibiting Hsp70 rebinding is essential for Glucocorticoid Receptor activity

**DOI:** 10.1101/2020.05.03.075523

**Authors:** Elaine Kirschke, Zygy Roe-Zurz, Chari Noddings, David Agard

**Affiliations:** Department of Biochemistry and Biophysics, University of California, San Francisco, CA 94158, USA

**Keywords:** Glucocorticoid Receptor, Hsp90, Hsp70, Bag-1, nucleotide exchange factor

## Abstract

The glucocorticoid receptor (GR), like many signaling proteins requires Hsp90 for sustained activity. Previous biochemical studies revealed that the requirement for Hsp90 is explained by its ability to reverse Hsp70-mediated inactivation of GR through a complex process requiring both cochaperones and Hsp90 ATP hydrolysis. How ATP hydrolysis on Hsp90 enables GR reactivation is unknown. The canonical mechanism of client release from Hsp70 requires ADP:ATP exchange, which is normally rate limiting. Here we show that independent of ATP hydrolysis, Hsp90 acts as an Hsp70 nucleotide exchange factor (NEF) to accelerate ADP dissociation, likely coordinating GR transfer from Hsp70 to Hsp90. As Bag-1 is a canonical Hsp70 NEF that can also reactivate Hsp70:GR, the impact of these two NEFs was compared. Simple acceleration of Hsp70:GR release was insufficient for GR reactivation as Hsp70 rapidly re-binds and re-inactivates GR. Instead, inhibition of GR re-inactivation by Hsp70 is critical. This can be accomplished by high non-physiological Bag-1 concentrations, which also inhibit Hsp70:ATP binding. In contrast, in an ATP-hydrolysis dependent process, Hsp90 plays a unique role by kinetically partitioning GR into a state that can bind ligand, but is protected from Hsp70 inactivation, thus allowing GR to be activated by its ligand but still able to re-enter the chaperone cycle. At physiologic concentrations, Bag-1 works synergistically with Hsp90 to accelerate the first rate-limiting step in GR reactivation. The net effect is that the chaperone machinery cyclically dictates the on and off rates for GR ligand, providing a timer controlling the persistence of activated GR.

**Significance Statement:** The glucocorticoid receptor (GR) is an essential transcription regulatory factor. Like many signaling proteins, GR activity is regulated by two essential molecular chaperones: Hsp90 and Hsp70. Functioning like a toggle switch, Hsp70 first inactivates GR, and then Hsp90 reactivates GR in an Hsp90 ATP hydrolysis dependent manner. Here, an intricate set of biochemistry experiments uncover fundamental principles governing how these chaperone systems collaboratively regulate GR activity. While Hsp90 promotes GR release from Hsp70 by modulating Hsp70’s nucleotide state, this occurs independently of Hsp90 ATP hydrolysis. Instead, ATP hydrolysis on Hsp90 facilitates a second essential reactivation step resulting in an Hsp90-bound GR state that protects GR from Hsp70 re-inactivation. A kinetic partitioning model best describes chaperone modulation of GR’s activity.

## INTRODUCTION

Hsp70 and Hsp90 are ubiquitous and highly conserved molecular chaperones that function as core components of protein folding machines. While utilized for enhanced stability during *de novo* folding and moments of stress, a select group of proteins continue to rely on chaperones under physiological conditions for continued activity. For Hsp90, these proteins are referred to as “client” proteins and are typically identified by their loss of activity and/or degradation in response to inhibition of ATP hydrolysis on Hsp90 (1, 2). Hsp90’s clients are numerous, diverse, and enriched in signaling proteins, making Hsp90 an essential hub in eukaryotic protein homeostasis (3, 4). The glucocorticoid receptor (GR), is one of the most well-known Hsp90 clients, and serves as a paradigm for Hsp90 client interactions. GR is an essential transcription regulatory factor. By directly binding DNA and nucleating regulatory complexes at particular sites in the genome, GR precisely controls the transcription of a broad set of distinct gene networks in different cellular contexts (5). In this way, GR governs many essential developmental and physiological processes (6). GR’s nuclear localization, DNA binding, and transcription regulatory activities all require glucocorticoid binding, either by its endogenous cognate hormone (cortisol in humans) or synthetic glucocorticoids such as dexamethasone (dex). In an ATP dependent manner, Hsp90 is essential for GR to bind ligand and become active (7, 8). While chaperone mediated GR activation has been investigated for decades, many fundamental aspects of this process remain unclear.

Early pioneering work utilizing crude reconstitution systems uncovered an entire system of molecular chaperones involved in steroid receptor activation that in addition to Hsp90 includes Hsp70 and a specific set of Hsp70 and Hsp90 cochaperones: Hsp40, Hop, and p23 (9–11). Contrary to *in vivo* behavior, rigorous *in vitro* investigations found that GR’s ligand binding domain (GRLBD) binds ligand without aid or enhancement from Hsp90 (12, 13). By contrast, rather than promoting GR activity, Hsp70 (with its Hsp40 cochaperone) was found to inhibit GR ligand binding by partially unfolding the GRLBD. Addition of Hsp90 (with Hop and p23) then completely reversed the Hsp70 inhibition (13). Furthermore, in the presence of the entire chaperone system, the affinity and rates for GRLBD ligand binding were enhanced. Importantly, the reversal of the Hsp70 inhibition requires ATP hydrolysis on Hsp90. Without Hsp90 ATP hydrolysis, GR remains in an inactive state bound to a stalled multi-chaperone complex containing both Hsp70 and Hsp90 (8, 13). This suggests that the Hsp90 dependence largely results from the necessity to reverse the Hsp70 inhibition through an intimate client hand-off and refolding process that requires ATP hydrolysis on Hsp90. Consistently, a synergistic collaboration between Hsp90 and Hsp70 in firefly luciferase refolding has since been observed for the bacterial, yeast, and human chaperone systems, further indicating a fundamental and universally conserved interconnection of the chaperone cycles (14–17).

Hsp70 and Hsp90 typically act at distinct phases in the folding pathway. Hsp70 recognizes fully extended hydrophobic polypeptide stretches and thus preferentially interacts with unfolded proteins (18–20). However, flexibility in Hsp70’s substrate binding domain (SBD) lid provides some degree of plasticity for binding exposed locally unfolded regions on otherwise structured proteins (21). This provides a rationale for how Hsp70 interacts with some near native proteins, and how GRLBD can exhibit only partial structural disruption while bound to Hsp70 (13, 22). Hsp70 substrate binding and release is regulated by nucleotide dependent conformational changes (23). When bound to ATP, Hsp70’s nucleotide binding domain (NBD) docks onto the SBD. NBD:SBD association stabilizes a state in which the SBD’s lid is open and the substrate binding pocket has switched to a low affinity state, allowing for substrate release (24). Opposingly, upon ATP hydrolysis the NBD dissociates from the SBD, allowing the SBD’s substrate binding site and lid to close, resulting in high affinity substrate binding (25, 26).

Hsp70 activity is heavily regulated by context specific co-chaperones, which typically function optimally at sub-stoichiometric ratios to allosterically modulate Hsp70 at specific states in its hydrolysis cycle to drive substrate binding and release in distinct biological and cellular contexts. Substrate loading is facilitated by a class of proteins referred to as J-domain containing proteins (JDP). JDPs assist substrate protein binding and via their conserved J-domain, stimulate ATP hydrolysis on Hsp70, leading to the tightly bound state (27). Hsp40, which is required for Hsp70 inhibition of GR, is a JDP. On the other hand, nucleotide exchange factors (NEFs) compensate for Hsp70’s slow ADP (and substrate) release rates by stabilizing a low nucleotide affinity NBD state (28), allowing ATP to rebind and substrate to release, and thus reset the cycle. Functioning equivalently to the bacterial NEF GrpE, the evolutionary distinct group of Bag domain containing proteins function as NEFs in the human system. While structurally distinct, both GrpE and Bag domains bind to the same surface of Hsp70 NBD and induce a low nucleotide affinity sate of the NBD (29). Although not as large and diverse as the JDP family, Bag domain containing proteins also possess additional functional domains that couple Hsp70 function to other cellular systems.

Hsp90 preferentially interacts with partially folded intermediate states populated later in the folding pathway (30). Hsp90 appears to have particularly complex client recognition principles that are still not well understood. This is further complicated by the complex ATP hydrolysis cycle in which Hsp90 undergoes dramatic conformational changes along a multistate cycle (31). In brief, Hsp90 functions as a homodimer with a high affinity dimerization interface at its C-terminal domain (32). In the absence of nucleotide Hsp90 forms a flexible V-shaped dimer (33, 34). ATP binding to the N-terminal domain promotes a secondary dimerization interface between the N-terminal domains which stabilizes a “closed”, hydrolysis competent Hsp90 conformation (35). Eukaryotic Hsp90 function is also heavy regulated by context specific co-chaperones that stabilize specific functional states along the Hsp90 hydrolysis cycle. The two co-chaperones required for GR activation, Hop and p23, are well established Hsp90 co-chaperones. Hop (*Hsp*70/Hsp90-Organizing Protein) facilitates client transfer from Hsp70 to Hsp90 by binding both Hsp70 and Hsp90 and stabilizing a conformational state of Hsp90 that appears preorganized for client binding and dimerization of Hsp90’s N-terminal domains (36–38). Acting last and mutually exclusive to Hop, p23 stabilizes the closed state of Hsp90 that is associated with GR activation (13, 35, 39).

Consistent with recognition of partially folded intermediates, Hsp90 has the capacity to bind substrates through both structured and unstructured client regions (40–42). Surprisingly, for all direct client interactions investigated *in vitro*, nucleotide only biases Hsp90’s confirmation, resulting in only minor-to-modest impact on client interaction affinities (12, 41, 43). Thus, a central unanswered question is how ATP binding and hydrolysis on Hsp90 are coupled to client folding and activation. It is now clear that ATP binding and hydrolysis on Hsp90 is not solely responsible for regulating direct client interactions, but is essential for coordinating an intimate functional coupling with Hsp70. Mirroring GRLBD ligand binding inhibition, Hsp70’s strong affinity for hydrophobic stretches forming the hydrophobic core of proteins appears to over stabilize unfolded protein states. This results in inhibition of protein folding at high, albeit physiological Hsp70 concentrations (16). Importantly, Hsp90 releases the inhibitory effect of Hsp70 and allows the client to fold at a spontaneous rate. For both protein refolding and GR ligand binding regulation, ATP hydrolysis on Hsp90 is essential for releasing Hsp70’s inhibitory effect. Exactly how ATP hydrolysis on Hsp90 releases Hsp70’s inhibition, directly through Hsp70 and/or clients is unclear.

Inspired by a direct contact between Hsp90 and Hsp70’s nucleotide binding domains observed in a low resolution cryo-EM structure of a GRLBD bound Hsp90:Hop:Hsp70 complex, we previously hypothesized that ATP hydrolysis on Hsp90 may drive ADP release on Hsp70 (13). This would enable Hsp90 to promote client release from Hsp70 in a functionally equivalent way to NEFs and coordinate client transfer within an Hsp70:Hsp90:Hop:GR complex. Supporting this, equivalent concentrations of the NEF Bag-1, a potent catalyst of Hsp70:ADP dissociation found associated with GR chaperone complexes, similarly reverse the Hsp70 induced inhibition of GRLBD (13, 44, 45). Here, the hypothesized NEF activity of Hsp90 is directly tested. While Hsp90 accelerates ADP:Hsp70 dissociation, it surprisingly does so entirely independent of ATP hydrolysis on Hsp90. This indicates that ATP hydrolysis on Hsp90 functions on another essential step in the GRLBD reactivation pathway other than Hsp70 nucleotide exchange. To gain insight into how ATP hydrolysis on Hsp90 promotes GR activation, we explored GRLBD chaperone regulation in deeper biochemical detail.

## RESULTS

### Hsp90 promotes Hsp70 nucleotide release independent of ATP hydrolysis on Hsp90

To determine if Hsp90 promotes GRLBD reactivation by promoting ADP release on Hsp70, the effect of Hsp90 on the dissociation rate of fluorescently labeled nucleotide from Hsp70 by fluorescence polarization was measured. This experiment was carried out in the presence of Hop, which stabilizes a functional interaction between the chaperones (38, 46). Furthermore, in the context of GR, Hop is required for full GRLBD reactivation, and stabilizes the N-terminal domain rotated state of Hsp90 required for the direct interaction between Hsp70’s and Hsp90’s ATPase domains observed in a cryo-EM reconstruction of the Hsp90:Hop:Hsp70:GR complex (13, 36). First, GRLBD, Hsp70, Hsp40, and ATP-FAM were pre-incubated to form the GRLBD:Hsp70:ADP-FAM complex. Dissociation of the fluorescently labeled nucleotide was then initiated by addition of excess unlabeled ATP alone, with stoichiometric amounts of pre-formed Hsp90:Hop complex, or Bag-1 as a positive control. Nucleotide dissociation was measured in the presence of 10mM phosphate, which slows ADP:Hsp70 dissociation, allowing the ADP dissociation rate to be captured without a stopped flow instrument (44). Remarkably, an ~3-fold acceleration in the nucleotide dissociation rate was observed with Hsp90:Hop (Fig.1). This acceleration is significantly less than the enhancement observed with an equivalent concentration of Bag-1, for which the nucleotide dissociation rate was too fast to measure. While experiments were carried out with GRLBD, GRLBD is not necessary for Hsp90 induced acceleration (Fig. S1). This is consistent with the formation of the Hsp90:Hop:Hsp70 complex in the absence of client (36).

**Fig. 1.**
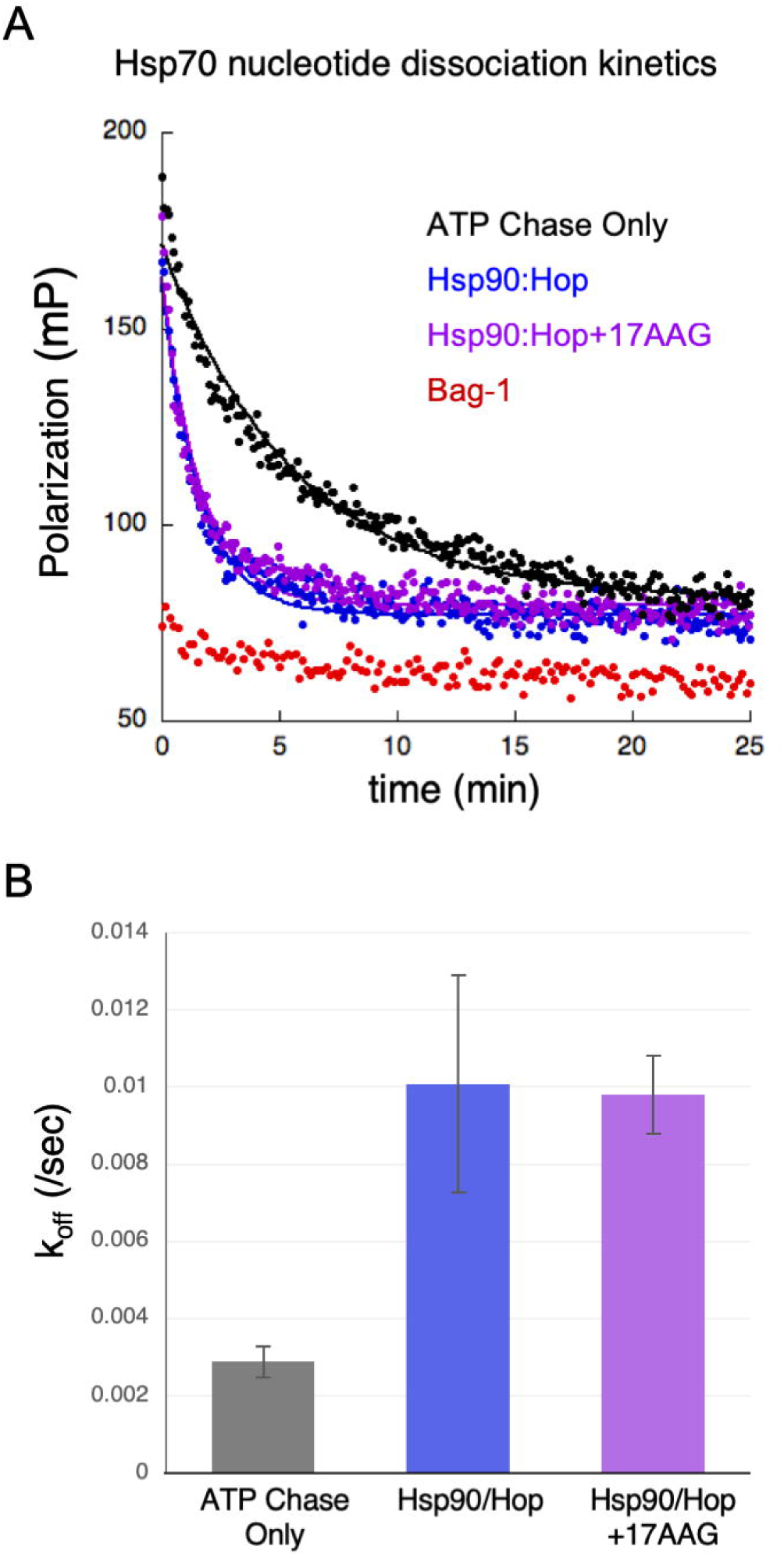
Hsp90 accelerates Hsp70 nucleotide dissociation. (*A*) 10μM GRLBD, 200nM ATP-FAM, 10μM Hsp70, and 1μM Hsp40 were pre-incubated to form the GRLBD bound Hsp70-ADP state. Kinetics were initiated by addition of 1mM ATP alone (black), or with 10μM Bag-1 (red), or 10μM Hsp90(dimer)/Hop, without (blue) or with 50µM 17AAG (purple). Curves were fit to a single exponential decay. (*B*) Average off rate as in A from three independent experiments with error bars representing S.E.M.. Average off rates are: 0.29±0.04 x10^3^ sec^−1^ for Hsp70 alone, 1.01±0.28 x10^3^ sec^−1^ for Hsp90/Hop, and 0.98±0.10 × 10^3^ sec^−1^ for Hsp90/Hop with 17AAG (±S.E.M.).

Surprisingly, acceleration of ADP dissociation by Hsp90 was not inhibited by the specific Hsp90 inhibitor 17AAG, which competes with ATP for Hsp90’s nucleotide binding pocket and inhibits ATP hydrolysis (Fig. 1). Similar results were obtained with Hsp90 (D93N), an ATP binding deficient mutant (90D) (Fig. S1). Together, this indicates that while Hsp90 promotes nucleotide release on Hsp70 to promote and consolidated Hsp70:Hsp90 client transfer, it does so independent of Hsp90 ATP binding and hydrolysis. Since ATP hydrolysis on Hsp90 is essential to reverse Hsp70’s inhibition of GRLBD, ATP hydrolysis on Hsp90 is required for an alternative step other than nucleotide release on Hsp70 that is essential for Hsp90 dependent GRLBD reactivation.

### Bag-1 Reverses Hsp70 Induced GRLBD Inhibition by Inhibiting Hsp70 ATP Rebinding

Given that both Hsp90 and Bag-1 reverse the Hsp70 inactivation of GRLBD, the requirement for a second essential step other than nucleotide exchange on Hsp70 for GRLBD activation by Hsp90 is surprising. In light of this, the mechanisms by which Bag-1 and Hsp90 reverse the Hsp70 inhibition of GRLBD were more carefully compared. Utilizing an *in vitro* reconstituted system in which GRLBD ligand binding to fluorescein labeled dexamethasone (F-dex) is monitored by fluorescence polarization anisotropy, addition of Bag-1 to Hsp70 pre-inhibited GRLBD was previously shown to result in rapid GRLBD reactivation (13). In this case, GRLBD reactivation correlates with the loss of GRLBD:Hsp70 binding. At the high concentrations of Bag-1 explored previously (stoichiometric to Hsp70), the rapid reactivation of GRLBD is followed by a gradual loss of GRLBD function. The slow loss of function appears to be due to aggregation of the partially unfolded GRLBD released from Hsp70. Here, using the same GRLBD ligand binding assay, Bag-1 was titrated to determine Bag-1’s effective concentration range for reactivating GRLBD (Fig. 2). The expectation was that Bag-1 would function at catalytic concentrations as it does for stimulating Hsp70 ADP release, ATP hydrolysis, and protein refolding (44, 47). Focusing on earlier time points and using the peak of GRLBD reactivation to determine approximate equilibrium levels of GRLBD ligand binding activity, an EC_50_ of 2.0µM for Bag-1 was obtained (Fig. 2*C*).

**Fig. 2.**
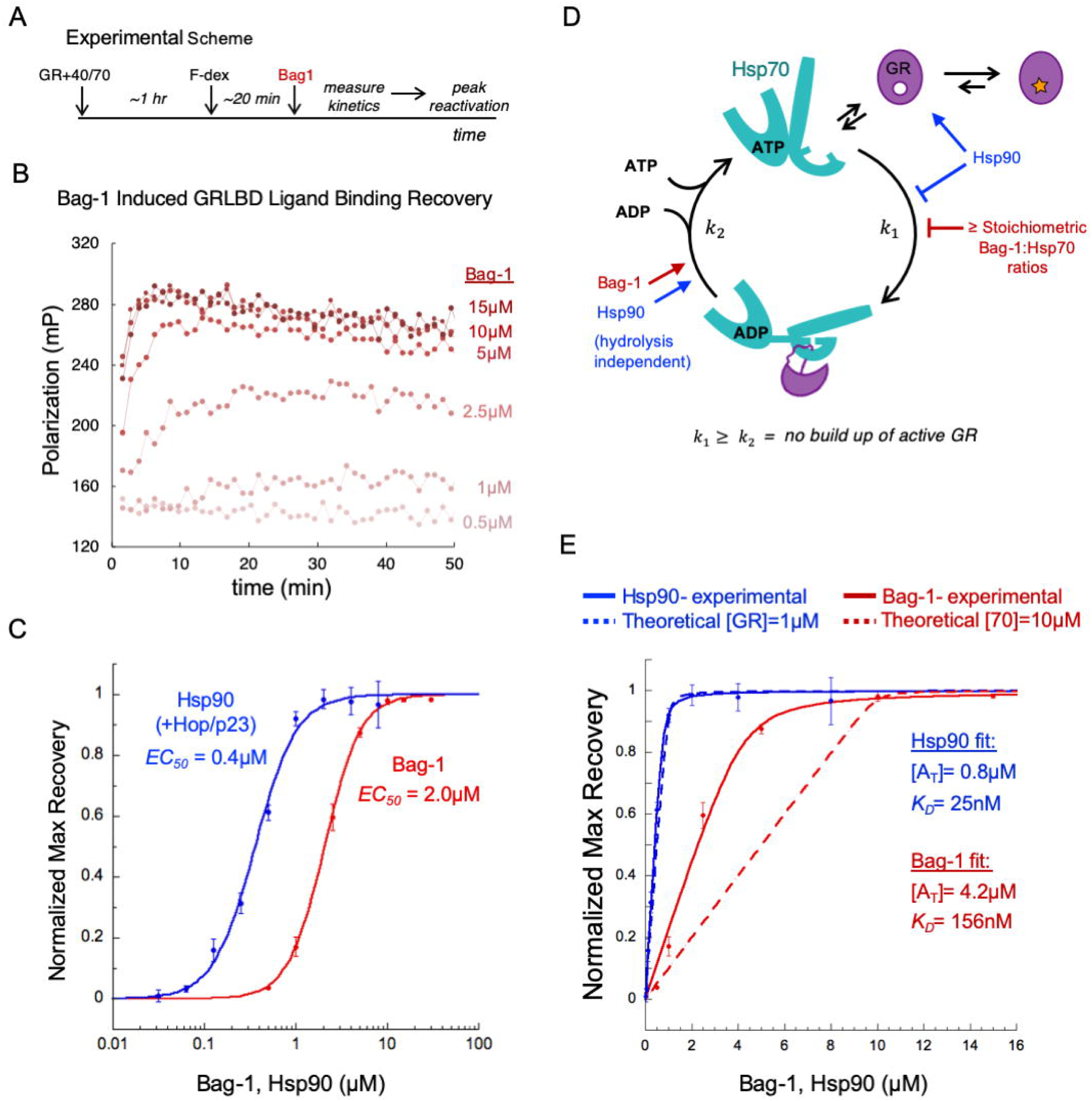
Equilibrium Bag-1 induced GRLBD ligand binding reactivation. (*A*) Experimental scheme. (*B*) Kinetics of Bag-1 induced GRLBD ligand binding reactivation. 1µM GRLBD was pre-inhibited by 10µM Hsp70 and 2µM Hsp40, and then equilibrated with 20nM F-dex. Reactivation kinetics were initiated by addition of varying concentrations of Bag-1. (*C*) The average maximum reactivation values of two independent experiments as in B are normalized and plotted against Bag-1 concentrations (red). Also shown is the equivalent reactivation with varying concentrations of Hsp90 dimer with 15µM Hop and p23 as reported previously (blue) (13). Error bars represent S.D.. Curves fit an EC_50_ of 2.04±0.00µM for Bag-1 and 0.37±0.02µM for Hsp90 (±S.E.M.). (*D*) Model for GRLBD inhibition and reactivation by Hsp70. E) Data from Fig. 2C plotted on a linear scale and fit with fraction bound equation for Hsp90 (solid blue line) and Bag-1 (solid red line). Dashed lines indicate theoretical fit (Fig. 3S) assuming GR reactivation through stoichiometric association of Hsp90_2_:GR (blue) or Hsp70:Bag-1 (red).

Comparing the effective concentration range for reactivation by Bag-1 to that of Hsp90 (with Hop and p23) under the same conditions, Hsp90 reactivated GRLBD at a ~5-fold lower concentration (EC_50_ = 0.37µM) (Fig. 2*C*). Due to the complexity of this multicomponent system, the pseudo-first-order approximation and thus the standard K_D_ equation do not apply. More specifically, under our reaction conditions GRLBD and Hsp70 were at 1 µM and 10µM respectively. While optimal for detection of GRLBD inhibition and reactivation, these concentrations are orders of magnitude higher than the nanomolar K_D_s for the titrated components (Bag-1 and Hsp90) (13, 47). Therefore, these plots more likely reflect stoichiometric properties of the system rather than true affinities, as in a stoichiometric saturation plot. It is hence noteworthy that the effective concentration range for reactivation by Bag-1 alone saturates much closer to Hsp70’s concentration (10µM), rather than the concentration of GRLBD (1µM). This indicates that as opposed to Hsp90 which acts stoichiometrically to GRLBD, Bag-1 acts more stoichiometrically to Hsp70 to promote GRLBD reactivation.

It is well appreciated that at sub-stoichiometric ratios, NEFs such as Bag-1 are potent catalysts of ADP release, resulting in acceleration of Hsp70’s steady state ATP hydrolysis levels when ADP release is rate limiting, such as in the presence of a JDP (44, 47). By contrast, high concentrations of Bag proteins at stoichiometric Bag:Hsp70 ratios inhibit steady-state ATP turnover (47). This is thought to result from the ability of Bag domains to preferentially bind and stabilize the apo state of Hsp70’s NBD, and in doing so, inhibit ATP binding. Consistent with previous reports, we directly observed the inhibition of Hsp70:ATP binding by Bag-1 at stoichiometric Bag-1:Hsp70 ratios (Fig S2). ATP hydrolysis on Hsp70 is essential for the inhibition of GRLBD ligand binding (13). Therefore, the equilibrium buildup of active GRLBD occurring at stoichiometric Bag-1:Hsp70 concentrations is largely due to inhibition of Hsp70:ATP rebinding. Together, this suggests a mechanism for Bag-1 reactivation in which Bag-1 rapidly stimulates GRLBD release from Hsp70 by promoting ADP dissociation, and then at higher, stoichiometric Hsp70:Bag concentrations, Bag-1 prevents Hsp70 re-inhibition of GRLBD by blocking the ability of Hsp70 to rebind nucleotide (Fig. 2*D*).

This model implies that at low Bag-1 concentrations, which catalytically promote Hsp70:ADP dissociation (47) but result in minimal GRLBD reactivation, any GRLBD released as a result of nucleotide exchange must be readily rebound and re-inhibited by Hsp70. The net result is cycling of GRLBD on and off Hsp70 without buildup of an active GRLBD population (Fig. 2*D*). Therefore, protecting GRLBD from inactivation by Hsp70 is critical for obtaining active receptor. Of note, the typical Bag-1 cellular concentration range (0.1-2µM) is 10 to 50 fold below that required to maintain an active GRLBD state through stoichiometric inhibition of Hsp70:ATP binding (7-22µM) (16, 48, 49). This indicates that steady state buildup of active GRLBD by Bag-1 alone is not likely physiological, and importantly, explains the cellular requirement for Hsp90 to maintain an active GR population despite the presence of Bag-1. Taken together, these results reveal that Hsp90 must do more than simply facilitate nucleotide exchange on Hsp70 to enable GRLBD reactivation. Since Hsp90’s effect titrates to GRLBD’s concentration as opposed to Hsp70’s, Hsp90 likely achieves GRLBD reactivation by directly binding GRLBD (12, 39) and, through an Hsp90:ATP hydrolysis dependent process, sequestering the GRLBD in an active state not readily re-inactivated by Hsp70.

Exploring this model further, we propose a simplified equilibrium based mechanism by which GRLBD reactivation is achieved through either Hsp90:GRLBD or Bag1:Hsp70 association (Fig S3), and fit the experimental GRLBD reactivation curves (Fig. 2*E*). This model describes Hsp90 induced reactivation remarkably well, suggesting that competitive inhibition of Hsp70 by Hsp90:GRLBD binding strongly accounts for GRLBD reactivation by Hsp90. However, stoichiometric ATP binding inhibition of Hsp70 by Bag-1 does not fully account for Bag-1 induced reactivation. It is unclear whether this is due to a more complex nucleotide displacement mechanism involving conformational changes on Hsp70 as suggested for GrpE (Fig. S4*C*) (50), preference of Bag-1 for interacting with Hsp70:ADP or Hsp70:ADP:GRLBD, or unaccounted kinetics effects related to nucleotide hydrolysis. Regardless of the mechanism, the net effect is that Bag-1 appears more potent than purely stoichiometric inhibition of Hsp70, but less potent than expected if only Hsp70 nucleotide exchange catalysis was required (Fig. S4).

### GRLBD Release from Hsp70 is Sufficient for Enhanced GRLBD Ligand Binding, but is Not Sustained by Bag-1

In the presence of the full chaperone system (Hsp70, Hsp90, Hsp40, Hop, and p23), GRLBD has a higher affinity for ligand, with an accelerated rate of both ligand association and dissociation (13). However, it was unclear to what extent the enhanced ligand binding ability was uniquely attained with Hsp90, or if it could equivalently be achieved with Bag-1. Since ligand binds to the core of the GRLBD, some degree of unfolding must be required for ligand entry and release. Supporting this, single-molecule force spectroscopy revealed that GRLBD’s Helix 1 must unfold for ligand to bind (51). We previously hypothesized that the enhanced ligand binding activity resulted from partial unfolding of the GRLBD such that the buried ligand binding pocket becomes accessible while interacting with the full chaperone system. Supporting this model, hydrogen-deuterium exchange measured by mass spectrometry previously demonstrated that interaction with Hsp70 induced partial un-structuring in specific regions of GRLBD that make direct contact with ligand (13). While providing an explanation for how Hsp70 inhibits GRLBD ligand binding, it also supports a mechanism in which Hsp70 “opens” the ligand binding pocket, such that upon Hsp70 release, the ligand binding pocket is transiently accessible. Here, we sought to clearly determine if GRLBD’s enhanced ligand binding ability is entirely due to release from Hsp70, and thus could be achieved with Bag-1, or to what extent additional action from Hsp90 is required.

To address this, GRLBD ligand association kinetics was measured either with the full chaperone system (Hsp40, Hsp70, Hsp90, Hop, and p23) or for Hsp70 (and Hsp40) inhibited GRLBD recently released by Bag-1 (concentrations stoichiometric to Hsp70) (Fig. 3). This was carried out at lower GRLBD concentrations in which the dex binding kinetics are relatively slow and easy to capture. Of note, for this GRLBD concentration regime the GRLBD:dex interaction is not saturating. As a result, any increase in the plateau value in the presence of the chaperone system reflects an increase in the GRLBD ligand affinity. Interestingly, the initial rate for GRLBD ligand association with Bag-1 was fast, matching that with the Hsp90 system. However, the increased plateau over GRLBD alone seen with the Hsp90 system is not observed with Bag-1 alone (Fig. 3*B*). These measurements were carried out for three GRLBD concentrations, and all the observed ligand association rates (k_obs_) with Bag-1 matched those obtained with the Hsp90 system (Fig. 3*C*). This indicates that GRLBD’s ligand on rate (k_on_) upon release from Hsp70 is the same in the context of both Bag-1 and Hsp90 (k_on_ ~0.3 µM^-1^min^-1^, compared to 0.1 µM^-1^min^-1^ for GRLBD alone). Therefore, the enhanced GRLBD ligand on rate is entirely due to GRLBD release from Hsp70, with no further action from Hsp90 required. However, the high equilibrium binding affinity in the presence Hsp90 suggests that Hsp90 is able to maintain this high ligand affinity GRLBD state for a longer period of time before it is ultimately released to then recycle back on Hsp70. This enables the enhanced ligand binding activity to be maintained through continual cycling with the chaperone system in a way that cannot be achieved with Bag-1 alone.

**Fig. 3.**
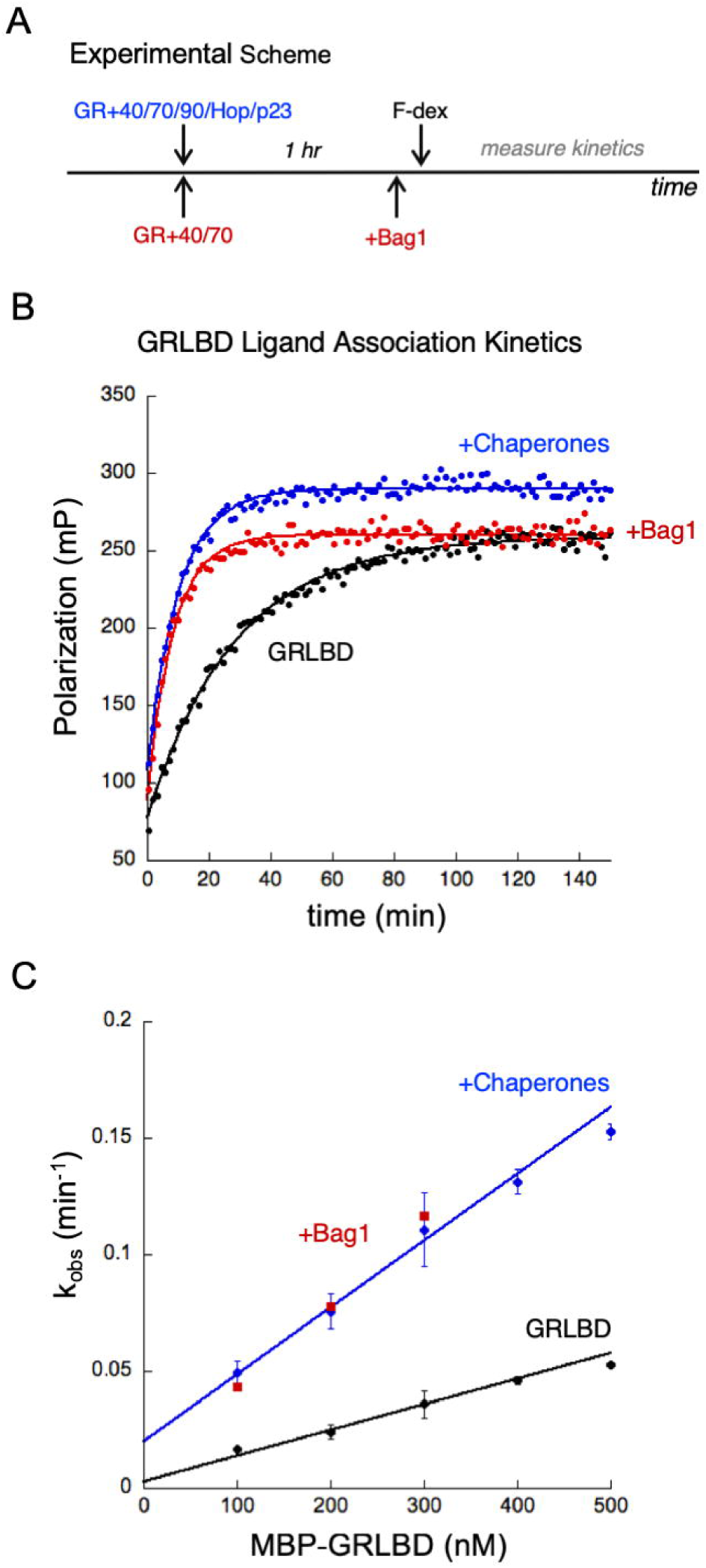
GRLBD ligand association kinetics. (*A*) Experimental scheme in which GRLBD was either pre-incubated with the full chaperone system (Hsp40, Hsp70, Hsp90. Hop, and p23) or with just Hsp40 and Hsp70 for ~1 hour. About 10 minutes before measuring GRLBD ligand association kinetics, Bag-1 was added to the Hsp40 and Hsp70 reaction to initiate GRLBD release from Hsp70. (*B*) Ligand association kinetics for 20nM F-dex binding to 300nM GRLBD either alone, or pre-incubated either with the entire chaperone system (2µM Hsp40, 15µM Hsp70, and 10µM Hsp90, Hop, and p23) or (2µM Hsp40, 15µM Hsp70, and 15µM Bag-1). (*C*) GRLBD ligand association rates (k_obs_) as in B, plotted against GRLBD concentration for GRLBD alone (black), with the full chaperone system (blue), and GRLBD released from Hsp70 by Bag-1 (red). For GRLBD alone and with chaperones, k_obs_ are the average of 3 independent experiments (± S.D.). Linear fit resulted in slopes corresponding to the ligand on rate (k_on_) of 0.11 µM^-1^min^-1^ for GRLBD alone and 0.29 µM^-1^min^-1^ with chaperones.

### Bag-1 Functions Synergistically with Hsp90 to Facilitate GRLBD Reactivation

While the kinetics of Bag-1 induced GRLBD reactivation are fast, the Hsp90 induced rate of reactivation is slow and exhibits a lag phase of about 10 minutes (13). This lag phase likely represents a rate-limiting step in a multistep client transfer process from Hsp70 to Hsp90 (Hsp90 ATP hydrolysis, client transfer, Hsp70/Hop release, Hsp90 closure). In the presence of both the Hsp90 system and high concentrations of Bag-1, the fast Bag-1 rate dominates (13). To determine if Bag-1 facilitates Hsp70 release and GR transfer in the Hsp90 pathway, Bag-1 was titrated in the presence of the Hsp90 system (with Hop and p23), and the kinetics of GRLBD ligand binding reactivation were measured (Fig 4). Intriguingly, Bag-1 reduced the lag phase at low, sub-stoichiometric Bag:Hsp70 concentrations (Fig. 4*B*).

**Fig. 4.**
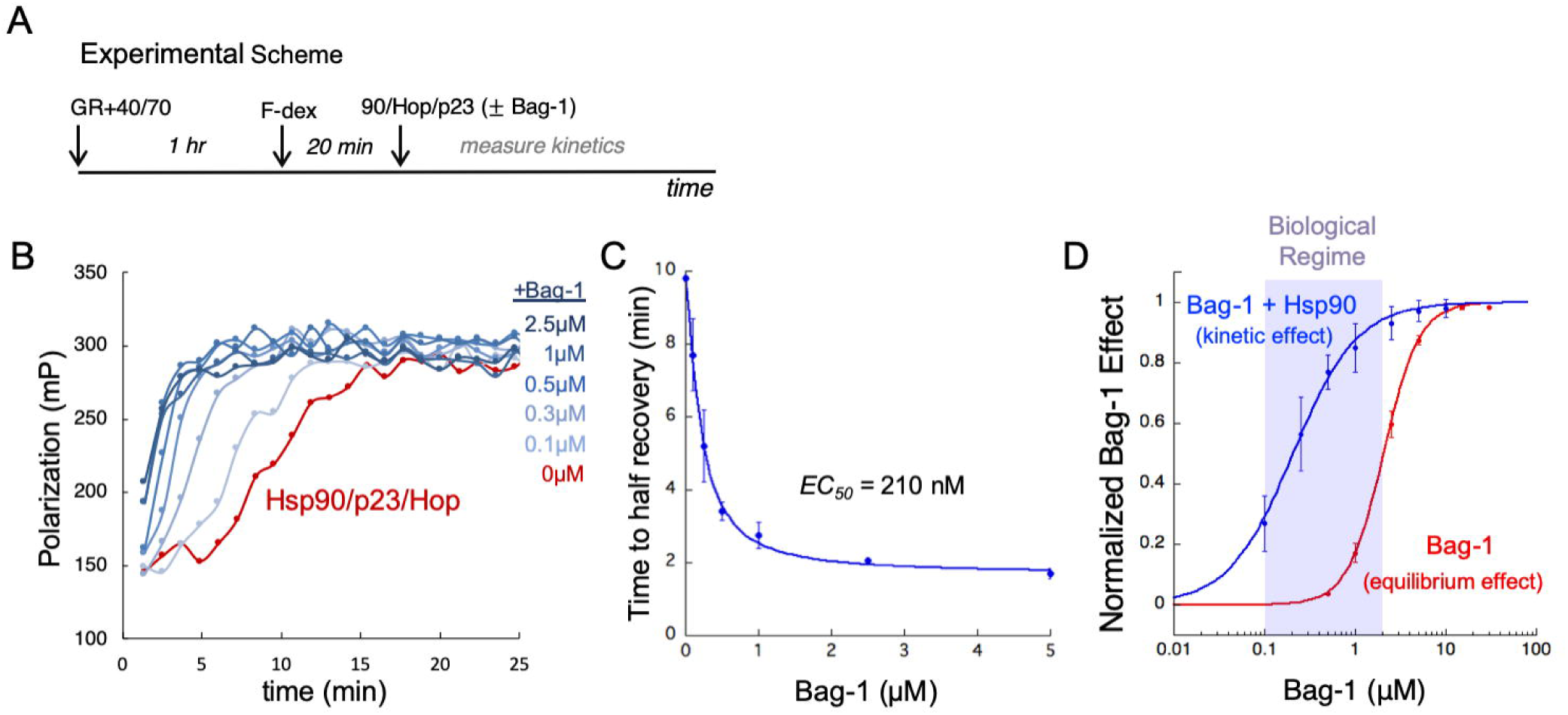
GRLBD reactivation kinetics (*A*) Experimental scheme. (*B*) 1µM GRLBD was pre-inhibited with 2µM Hsp40 and 15µM Hsp70. Ligand binding reactivation was initiated with 10µM Hsp90, Hop, and p23 premixed with varying concentrations of Bag-1 (0-2.5µM). (*C*) The average half time to recovery from two independent experiments was plotted against the corresponding Bag-1 concentration (± S.D.). Curve fit an EC_50_ of 0.21±0.02µM (±S.E.M.). (*D*) Normalized Bag-1 effect for Bag-1 GRLBD equilibrium reactivation as in Fig. 2*C* (red) and for Bag-1 acting with Hsp90 system to accelerate reactivation rate (blue). Biological Bag-1 concentration regime (0.1-2µM) indicated in purple.

Using the time to half recovery as a metric for GRLBD reactivation rate, an EC_50_ of 0.21µM for Bag-1 was obtained, with the effect saturating at a ~5-fold shorter halftime (Fig. 4*C*). Thus, for the same Hsp70 and GRLBD concentrations, the kinetic effect of Bag-1 with the Hsp90 system occurs at a 10-fold lower concentration of Bag-1 than does the equilibrium effect observed when Bag-1 acts independently. To clearly illustrate this, we plot together the normalized Bag-1 equilibrium effect for Bag-1 alone as in Fig. 2*C* and the kinetic effect seen in the context of the Hsp90 system (Fig. 4*D*). This demonstrates that Bag-1 has a pronounced ability to accelerate the rate of GRLBD reactivation at sub-stoichiometric Bag:Hsp70 concentrations for which no equilibrium GRLBD reactivation is observed by Bag-1 alone. This further supports the model in which sub-stoichiometric ratios of Bag-1 catalytically promote GRLBD release by ADP:ATP exchange but no GRLBD activity is detected due to its rapid re-inactivation by Hsp70. However, in the presence of the Hsp90 system, presumably due to Hsp90’s ability to trap GRLBD and inhibit Hsp70:GRLBD rebinding, the highly efficient Bag-1 NEF effect becomes pronounced. Given that Bag-1 is a significantly more potent catalyst of Hsp70:ADP dissociation than Hsp90, the rate limiting step in the Hsp70:Hsp90 GRLBD transfer resulting in the lag phase in the absence of Bag-1 is likely Hsp70:ADP dissociation. In this way, Bag-1 functions within the Hsp90 pathway at catalytic concentrations to accelerate the rate-limiting step for Hsp90 dependent GRLBD reactivation, resulting in more efficient GRLBD transfer and activation. Of note, the kinetic effect observed with the Hsp90 system occurs well within Bag-1’s cellular concentration regime (Fig. 4*D*), suggesting this more likely represents the physiological mechanism by which Bag-1 normally functions within the GR pathway.

Since ATP hydrolysis on Hsp90 is essential to reverse the Hsp70 inactivation of GRLBD, investigating the effect of Bag-1 in the context of hydrolysis deficient Hsp90 mutants enables the functional synergism between Bag-1 and the Hsp90 system to be studied without background reactivation from Hsp90. There are two well characterized Hsp90 ATP hydrolysis deficient mutants: D83N (Hsp90(D)), which cannot bind ATP, and hence blocking the formation of the ATP stabilized Hsp90 closed state, and E47A (Hsp90(E)), which can bind ATP and access the closed ATP stabilized state to which p23 binds but cannot hydrolyze ATP (Fig. 5*A*) (52, 53). In the absence of Bag-1, neither mutant reactivated GRLBD (13). Utilizing these Hsp90 mutants, the Bag-1 induced GRLBD reactivation assay described in Fig. 2 was carried out for varying concentrations of Bag-1 in the presence of the Hsp90 system (with Hop and p23). Intriguingly, an enhancement of the Bag-1 effect is observed in the presence of the hydrolysis deficient Hsp90 system, with GRLBD reactivation detected for lower Bag-1 concentrations compared to Bag-1 alone (Fig. 5*B*).

**Fig. 5.**
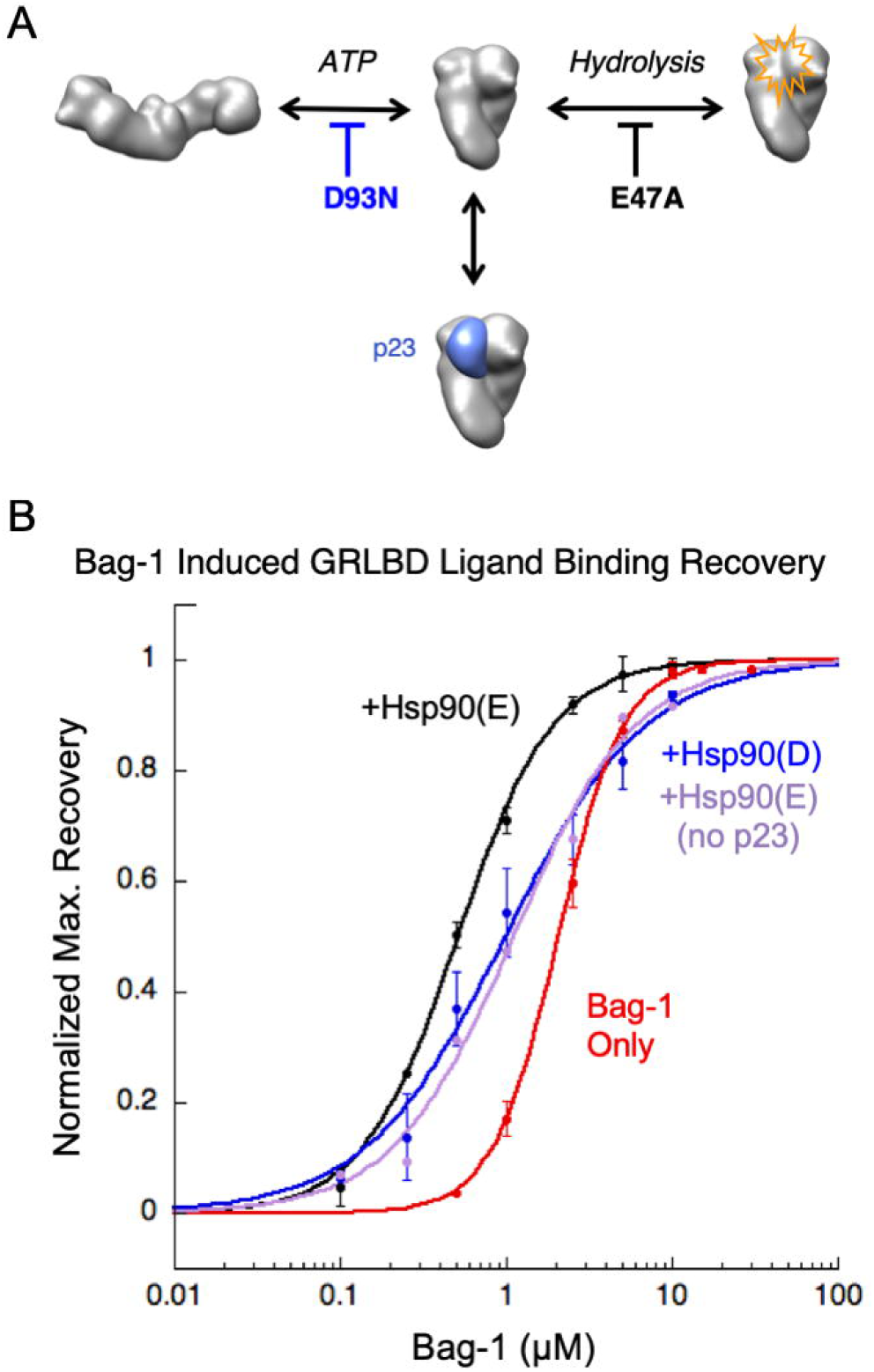
Hydrolysis dead Hsp90 enhances Bag-1 dependent GRLBD reactivation. (*A*) In the absence of ATP, Hsp90 dimers form a flexible V-shape, dimerized through the C-terminal domains. ATP binding to the N-terminal domains promotes formation of a second dimerization interface that stabilizes the closed state of Hsp90, to which p23 binds. Dimerization of the N-terminal nucleotide binding domains stabilizes Hsp90’s hydrolysis competent state. D93N prevents ATP binding, and thus inhibits Hsp90 closure. Hsp90 E47A can bind ATP, and thus close and bind p23, but cannot hydrolyze ATP. (*B*) Bag-1 induced GRLBD reactivation as in Fig. 2 (red), but with addition of the Hsp90 system (+Hop and p23) for either Hsp90 D93N (blue), Hsp90 E47A (black) and without (purple) p23. Data points represent the normalized average maximum reactivation values of two independent experiments (± S.D.). Data fit an EC_50_ of 2.04±0.00µM for Bag-1 alone, 1.03±0.21µM with Hsp90(D93N), and 0.52±0.00µM with Hsp90(E47A) (±S.E.M.).

Interestingly, the enhancement from the hydrolysis deficient Hsp90 system is more significant for the E47A mutant. To determine if the enhancement is due to ATP binding itself or the probability of forming the p23 stabilized closed state, the Bag-1 titration was also carried out with Hsp90(E47A) with Hop, but without p23. The absence of p23 with Hsp90(E47A) results in a trend matching that of Hsp90(D) (Fig. 5*B*). This indicates that the extra enhancement is due to the ability of Hsp90 to access the closed p23 bound state. Thus, even in the absence of ATP hydrolysis on Hsp90, the Hsp90 system provides a functional benefit in combination with Bag-1 to increase the equilibrium population of active GRLBD, with the strength of this effect dependent on Hsp90’s conformational state. A possible explanation for the enhancement resulting from the mutant Hsp90s is that even in the absence of ATP hydrolysis, Bag-1 can catalytically release client from Hsp70, after which Hsp70 and Hop dissociate from Hsp90. The data suggest that the ability of Hsp90 to sequester GRLBD, thereby blocking Hsp70 re-inhibition, correlates with the ability of Hsp90 to reach the closed state (stabilized by ATP and p23). Together, this supports a model in which Bag-1 and Hsp90 are functionally optimized for distinct and complimentary aspects of GRLBD reactivation such that they function synergistically to modulate GRLBD activity.

## DISCUSSION

Remarkably, Hsp90 promotes client release from Hsp70 by directly modulating Hsp70’s nucleotide state. This further supports a tight coupling of the Hsp70 and Hsp90 chaperone cycles during client transfer and growing evidence for a direct physical and functional interaction between Hsp70 and Hsp90 that is conserved across species (13–15, 54). However, the ability to do so independent of ATP binding and hydrolysis on Hsp90 indicates that Hsp90 ATP hydrolysis must promote GRLBD reactivation through another essential step in client transfer and activation other than ADP release from Hsp70. A careful comparison between GRLBD reactivation by Hsp90 and Bag-1 uncovered a previously unappreciated requirement for not just stimulating Hsp70 release, but also limiting re-inhibition by Hsp70. Both Bag-1 and Hsp90 facilitate client release by stimulating nucleotide exchange on Hsp70, although Bag-1 is far more potent at doing so. However, Bag-1 and Hsp90 achieve sustained GRLBD reactivation in the presence of Hsp70 through two distinct mechanisms that limit Hsp70:GRLBD re-inhibition. At stoichiometric concentrations, Bag-1 stimulates ADP release but also inhibits ATP rebinding to Hsp70, and consequently blocks the ability of Hsp70 to bind and inactivate GRLBD. However, this is unlikely to be biologically relevant due to the non-physiological Bag-1 concentrations required. In contrast, Hsp90 appears to prevent GRLBD re-inactivation by remaining associated with GRLBD after Hsp70 release and holding GRLBD in active state to which Hsp70 cannot rebind.

An important distinction between these two mechanisms emerges in regards to Hsp70 dependent enhancement of GRLBD ligand binding activity. Since Hsp70 release is sufficient to obtain the high ligand affinity GRLBD state, the enhanced ligand binding is entirely the result of “work” done by Hsp70 to make GRLBD’s ligand binding pocket accessible. As suggested by single molecule force experiments (51), this is likely achieved through displacement of GRLBD’s Helix1 during partial unfolding by Hsp70. A key difference between Hsp90 and Bag-1 is that while Bag-1 can transiently produce active GRLBD, this state is so short lived that no functional GRLBD can accumulate. Persistence of active GRLBD requires that Bag-1 inactivate the complete pool of Hsp70, which only happens at non-physiological Bag-1 concentrations. By contrast, Hsp90 functions to sequester the activated GRLBD in a manner that both allows ligand to bind and slows subsequent rounds of inactivation by Hsp70.

Support for this ability of Hsp90 to prolong the dwell time of an active partially unfolded client state generated by Hsp70, comes from recent experiments with the client protein Argonaut. There, Hsp70 enables Ago2 to partially populate a more extended Ago2 active conformation (55). This active state is then captured and stabilized by Hsp90. Consistencies between GR and Argonaut, two completely unrelated clients, suggest that this mode of combined Hsp70/Hsp90 chaperone action may apply to other constitutive Hsp90 client proteins that require Hsp90 throughout their functional lifetimes. This is likely particularly relevant in instances where access to a partially unfolded state not readily accessed under native conditions is beneficial for client activity.

Expanding upon this, it is becoming apparent that the key to understanding the functional effects of Hsp70 and Hsp90 lie in the kinetics of the chaperone cycles. Accordingly, we propose chaperone regulation of GR ligand binding activity can be explained by a kinetic partitioning model. In this model, GR’s observed ligand binding activity level is dictated by rate constants governing transitions along the chaperone cycle that determine the dwell times of each state in the pathway and therefore how GR partitions between free GRLBD (active, with pre-existing equilibrium between inactive and highly active states), Hsp70 bound GRLBD chaperone complexes (inactive), and Hsp70 free Hsp90 complexes (highly active) (Fig. 6). The resulting apparent GR ligand K_D_ is thus a net sum of the ligand affinities for the varying GRLBD conformations weighted by their population distribution, such that a longer dwell time in high affinity open ligand binding pocket state produces a higher apparent GR ligand affinity. In this light and consistent with early reports by Pratt and co-workers, the contribution of p23 to GRLBD’s Hsp90-dependent ligand binding activity can be explained by p23’s ability to bind to and prolong the dwell time of the closed nucleotide bound state of Hsp90 from which Hsp70 and Hop have dissociated. (Fig. 6) (39, 56). The newly appreciated significance is that the closed Hsp90 state can sequester GRLBD in its active conformation and that this is important to delay re-entry into the chaperone cycle, and thus Hsp70 re-inhibition. Generalizing beyond GR, this kinetic partitioning model aligns well with the report by Buchner and co-workers demonstrating that it is not Hsp90’s ATP rate (set by the rate limiting step in a multi-step ATP hydrolysis cycle) per se that matters, but instead the dwell time of Hsp90 in its different conformation states that is functionally important (57).

**Fig. 6.**
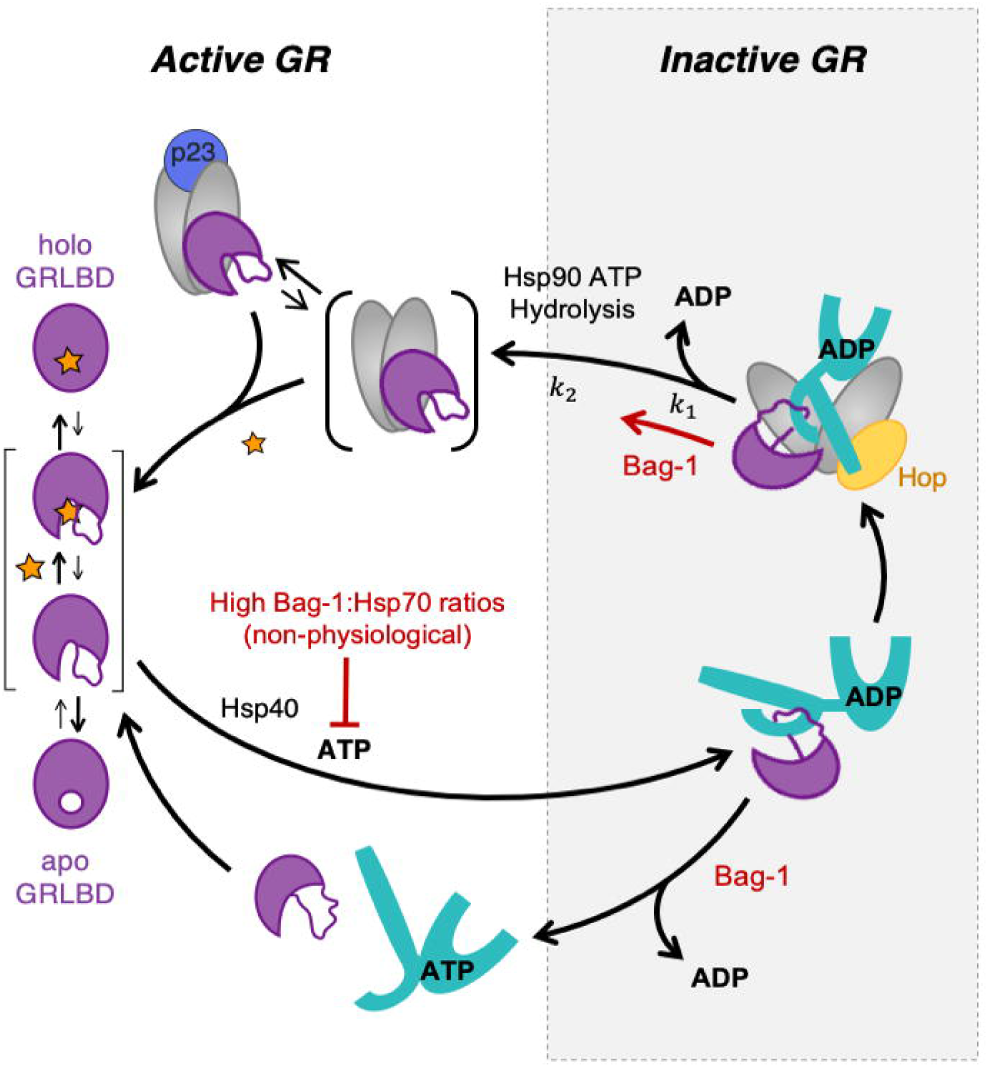
Kinetic partitioning model describes chaperone regulation of GR ligand binding activity. Kinetics rates governing transitions along the chaperone cycle bias how GRLBD partitions between inactive (Hsp70 bound) and active (Hsp70 free) states, with Hsp70 driving the opening of GRLBD’s ligand binding pocket to allow for enhanced ligand binding in the process. The net result is GR’s ligand binding activity level being set by the distribution of the GRLBD population between the different states with their corresponding ligand binding activities. Due to the rapid rate of Hsp70 re-inactivation, GRLBD activity in the presence of Hsp70 requires both Hsp70 release and inhibition of Hsp70 rebinding. Thus, GRLBD reactivation by Hsp90 requires at least two-steps, the first beings ADP dissociation on Hsp70 (Hsp90 ATP hydrolysis independent) followed by a second (Hsp90 ATP hydrolysis dependent) step that enables Hsp90 to sequester GRLBD in a high ligand affinity state that Hsp70 cannot bind. P23 helps maintain GRLBD activity by prolonging the dwell time of the high ligand affinity GRLBD state bound to the closed Hsp90 state. Bag-1 facilitates GRLBD release either from Hsp70:GRLBD or Hsp70:Hop:Hsp90:GRLBD complexes by accelerating ADP release from Hsp70, however buildup of an active GRLBD population without Hsp90 requires non-physiological stoichiometric Bag-1:Hsp70 concentrations that inhibit ATP rebinding to Hsp70 and thus GRLBD re-inactivation.

While little to no GRLBD activity is detected for low Bag-1:Hsp70 ratios in which Bag-1 catalytically promotes Hsp70 ADP release, likely due to the fast rate of Hsp70:GRLBD re-inactivation, the functional significance of Bag-1 becomes quite apparent when combined with the Hsp90 system where a dramatic reduction in lag phase is observed. Overall, our findings align remarkably well with early investigations by Pratt and co-workers and importantly provide a mechanistic explanation for the surprisingly opposing functional affects that high and low Bag-1:Hsp70 ratios had on chaperone-dependent GR activation by a reconstitution system (45). In this case, low ratios promoted progression of the chaperone cycles, while high Bag-1 ratios had a negative effect on GR’s chaperone dependent activity, which we show here is due to inhibition of entry into the chaperone cycle. Notably, Bag-1’s kinetic affect for facilitating client transfer occurs well within cellular Bag-1 concentration regimes, suggesting that client transfer to Hsp90 and other downstream systems, as opposed to promoting Hsp70 free GR, more accurately describes the physiological mechanisms by which Bag-1 functions. The degree to which this holds true for other client proteins depends on how rapidly they rebind Hsp70. Likely due to GRLBD’s dynamic nature, even in its “native” state GRLBD constantly accesses states that allow for rapid Hsp70 rebinding. By contrast, beyond initial folding which would utilize Hsp70, a more stable protein would attain a native state not accessible by Hsp70, effectively removing itself from a GR-like chaperone cycle. In such a case, NEFs could have positive functional effects at sub-stoichiometric concentrations. Such appears to be the case for Hsp70 dependent luciferase refolding in the absence of Hsp90, in which the NEF effect typically peaks at sub-stoichiometric NEF:Hsp70 ratios, before Hsp70 client binding inhibition at higher NEF:Hsp70 ratios begins to dominate (47). It is also worth noting that in the case where neither transfer to downstream chaperones nor a stable native state can be rapidly achieved, fast client recapture by Hsp70 would be desirable to protect clients from off pathway aggregation reactions. In collaboration with Hsp90, the intimate Hsp70:Hsp90 client hand off ensures minimal time spent dissociated from chaperones and provides the strongest anti-aggregation protection.

Once the inactive Hsp70:Hop:Hsp90:GRLBD complex is formed, generating active GRLBD requires transfer of client to Hsp90; dissociation of Hsp70 and Hop; closure of Hsp90 to form the active sequestered GRLBD. Since the rate limiting step resulting in the lag phase is nucleotide exchange on Hsp70, which Hsp90 accelerates independent of ATP hydrolysis, ATP hydrolysis on Hsp90 must function in a step post-nucleotide exchange. Several possible (non-mutually exclusive) mechanisms can be imagined where ATP hydrolysis on Hsp90 is required 1) to modulate the Hsp70 conformation within the Hsp70:Hop:Hsp90 complex to facilitate client transfer to Hsp90; 2) to act primarily on Hsp90 to dissociate Hsp70 and Hop from the complex; or 3) to drive a conformational change on the GRLBD, such as an unfolded to folded transition, that both correlates with GRLBD activation and occlusion of Hsp70 rebinding. Ultimately resolving this will require atomic structures of the various client bound chaperone states (Fig 6).

In conclusion, this work reveals that Hsp70:Hsp90 client hand off is even more complex and highly coordinated than previously appreciated. The tight coupling achieved through a multistep client hand off in which Hsp90 actively participates through direct Hsp70:Hsp90 contacts helps consolidate the chaperone cycles and ensures constant chaperoning throughout the client’s folding trajectory. Furthermore, an important biological advantage implied by the kinetic partitioning model is the potential for precise tuning and modulation of GR ligand binding activity level (as well as the activity level of other Hsp70/Hsp90 clients), particularly in different tissue and cellular contexts. This could be achieved either by specific auxiliary co-chaperones (Aha-1, FKBP51/52, and NEFs for example) and/or post translational modifications that regulate distinct steps in the chaperone cycle. Intriguingly, the kinetic partitioning model suggests that even in the presence of ligand, GR molecules continuously cycle between on and off states, which may have potentially interesting implications for GR’s transcription regulatory activities. Thus, the precise regulatory potential enabled by kinetic partitioning in the chaperone system further justifies the continued interaction of GR, as well of other clients, with this complex ATP consuming chaperone system.

## MATERIALS AND METHODS

### Protein Expression and Purification

Human Hsp90α, Hsp70 (HSPA1A), Hop, p23, the 26kDa Bag-1 isoform (116-345) and yeast Hsp40 (Ydj1) were expressed and purified as previously described (13). Human GRLBD (521-777) (F602S) was expressed with an N-terminal MBP tag in the presence of dexamethasone, which was removed by excess dialysis as previously described (13). The MBP tag previously shown to not interfere with the Hsp70 and Hsp90 chaperone cycle was maintained for enhanced GRLBD solubility. For simplicity, apo MBP-GRLBD(F602) is referred to as GRLBD.

### Fluorescence Polarization Measurements

All fluorescence polarization measurements were measured on SpectraMax M5 plate reader (Molecular Devices) with excitation/emission waves lengths of 485/538 nm and temperature control set to 25°C. Ligand binding reactions using fluorescein labeled dexamethasone (F-Dex) were carried out in 30mM HEPES pH 7.5, 150mM KCl, 5mM ATP, 5mM MgCl_2_, and 2mM DTT. Nucleotide dissociation reactions were carried out in 50mM HEPES pH7.6, 50mM KCl, 10mM Phosphate, and 5mM MgCl_2_ with N6-(6-Amino)hexyl-ATP-5-FAM (ATP-FAM).

### Hsp70 ADP Dissociation

10μM Hsp70, 10μM GRLBD, 1μM Hsp40 (Ydj1), and 200nM ATP-FAM were pre-incubated at room temperature for 5 minutes to obtain the ADP state of Hsp70. Dissociation kinetics were initiated by addition of 1mM ATP, either alone or with 10µM Hsp90_2_:Hop (±50µM 17AAG), or 10µM Bag-1. To promote formation of the Hsp90:Hop complex, Hsp90 and Hop were pre-incubated at room temperature for 5 minutes before combing with ATP. Curves were fit to a single exponential decay:

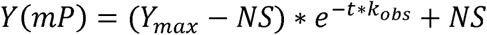

### GRLBD Equilibrium Ligand Binding Reactivation

1µM GRLBD was pre-incubated with 2µM Hsp40, 15µM Hsp70, and 5mM ATP/MgCl_2_ for 40-60 minutes at room temperature before equilibration with 50nM F-Dex at 25°C for about 20 minutes. F-Dex binding was initiated by addition of Bag-1 at varying concentrations. The average polarization value for the time window in which peak reactivation was observed was used as the approximate equilibrium value. The equilibrium data was fit using a cooperative binding model:

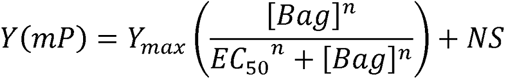

The slope (n=2.2±0.2 (S.E.M)) is not interpreted as meaningful due to concentration regimes. GRLBD reactivation by Hsp90 is previously described (13). Hsp90 equilibrium reactivation curve was fit using the same equation but substituting [Hsp90 dimer] for [Bag-1].

For analysis of the GRLBD ligand binding reactivation curves as equilibrium saturation plots, data was fit with the fraction bound equation for the standard reversible two component reaction mechanism derived in Fig. S3:

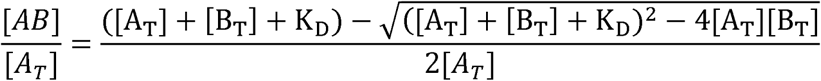

Where A = GRLBD and B = Hsp90, or A = Hsp70 and B = Bag-1.

For Bag-1 induced GRLBD ligand binding reactivation in the presence of hydrolysis dead Hsp90 system, reactions were carried out as for Bag-1 alone but with the addition of 10µM Hsp90 (E47A or D93N), Hop, and p23, which were pre-incubated in the presence of varying concentrations of Bag-1 for 30-60 minutes before addition with Bag-1 to Hsp70 inhibited GRLBD. Curves were fit as for Bag-1 alone for visual purposes only. Curve parameters were not interpreted due to the complexity of the reaction.

### GRLBD Ligand Association Kinetics

Varying concentration of GRLBD were pre-incubated either with the full chaperone system (2µM Hsp40, 15µM Hsp70, 10µM Hsp90,10µM Hop, 10µM p23) for about 60 minutes or 2µM Hsp40 and 15µM Hsp70 for about 50 minutes before addition of 15µM Bag-1 for 10 minutes. Incubations were carried out at room temperature in 5mM ATP and MgCl_2_. GRLBD ligand association kinetics were initiated by addition of 20nM F-dex to either MBP-GRLBD alone or Association curves were fit to a single exponential to determine the association rate (k_obs_):

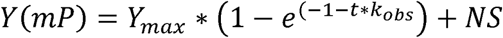

### Rate of Hsp90 Induced GRLBD Ligand Binding Reactivation with Bag-1

Experiments were carried out as described for the Bag-1 induced reactivation with the Hsp90 system but with wild type Hsp90. The average time to half recovery (t_1/2_) was used as an approximate reactivation rate, and plotted against the Bag-1 concentration to obtain an EC_50_ for Bag-1 using the following equation:

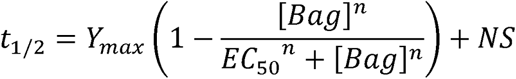

## Author contributions

Z.R.Z. carried out Hsp70 nucleotide dissociation experiments. E.K. carried out GRLBD ligand binding experiments and assembled all the data. E.K. and D.A.A. wrote the manuscript.

## ACKNOWLEDGMENTS

We thank Jason Gestwicki for the helpful discussions regarding development of the Hsp70 nucleotide dissociation experiments. This work was supported by the Howard Hughes Medical Institute (D.A.A.) and the National Science Foundation (Z.R.Z.).

---

**Fig. S1.**
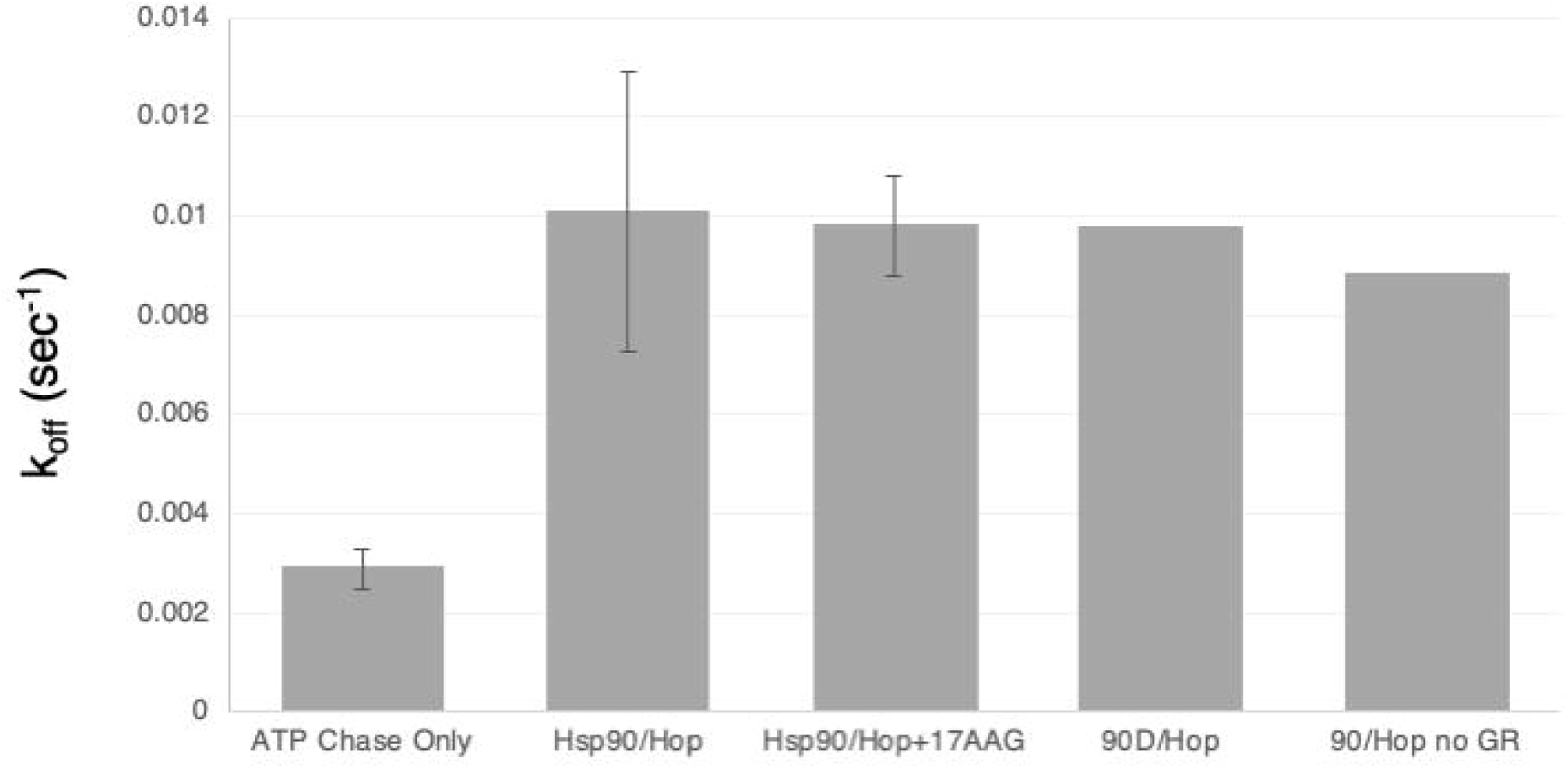
Acceleration in ADP release from Hsp70 by Hsp90 does not require ATP hydrolysis on Hsp90 or GRLBD. Rates of ADP dissociation as in Fig. 1*C*, but additionally including rates obtained with the hydrolysis deficient mutant of Hsp90 (D93N) and for wildtype Hsp90 and in the absence of GRLBD.

**Fig. S2.**
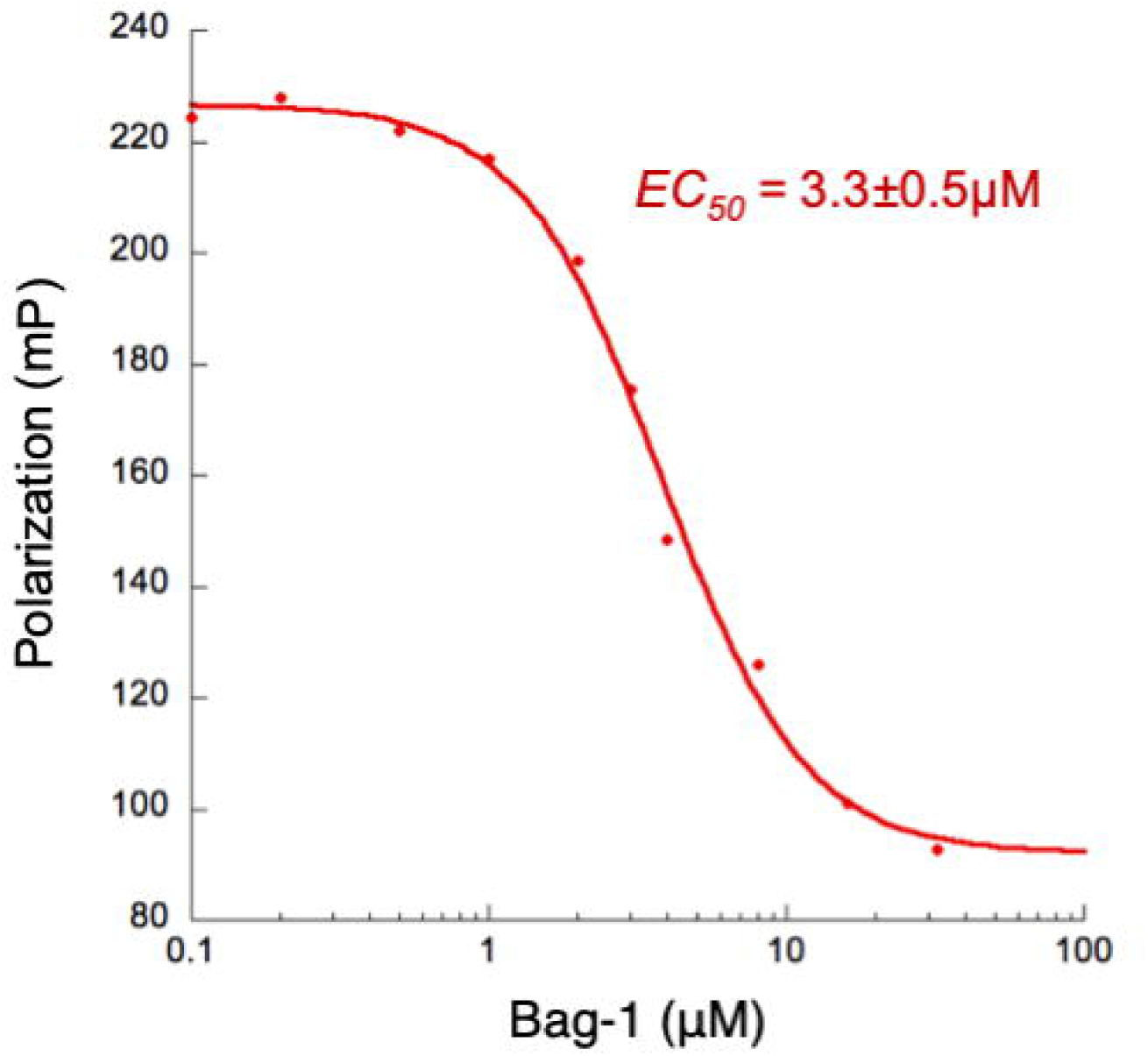
Stoichiometric concentrations of Bag-1 inhibit Hsp70 ATP binding. Equilibrium ATP binding to Hsp70 was measured by fluorescence polarization of 50nM ATP-FAM with 1µM Hsp70 and varying concentrations of Bag-1 in the same buffer conditions as for GRLBD ligand binding. A) Representative inhibition curve was fit using the following EC_50_ equation:

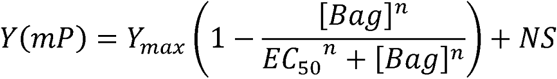

An average of two independent experiments yielded an EC_50_ of 3.3±0.5µM (SEM).

**Fig. S3.**
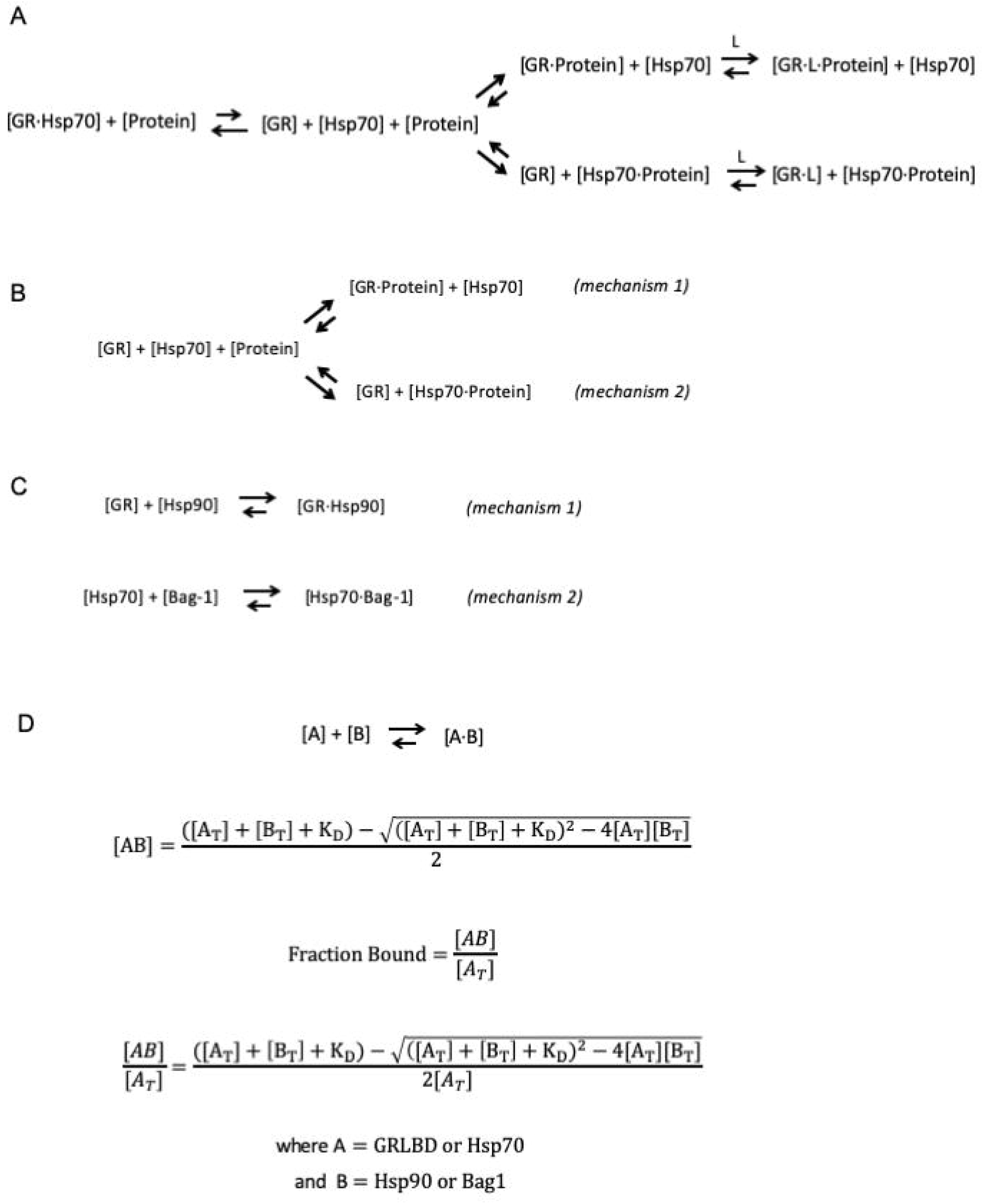
*A*) Mechanistic scheme depicting two different modes for reversing Hsp70 induced GRLBD ligand binding inhibition. (*B*) Simplified schematic from (A) in which certain reaction components are considered negligible due to experimental concentration regimes. In Fig. 2, the Hsp70 concentration is nearly saturating compared GRLBD, so we approximated that all free GRLBD will associate with Hsp70. Therefore Hsp70:GRLBD association was ignored. Since GRLBD is saturating compared to ligand (L), we approximated that all free GRLBD will bind ligand. Therefore, ligand binding was also ignored. (*C*) Further simplification of schematic from (B), leavening out nonreactive species for the proposed mechanism in the case of Hsp90 (top) and Bag-1 (bottom). D) The reaction schemes from (C) can be generalize to a two-component reversible association reaction in which the fraction bound can be determined by the following equation.

**Fig. S4.**
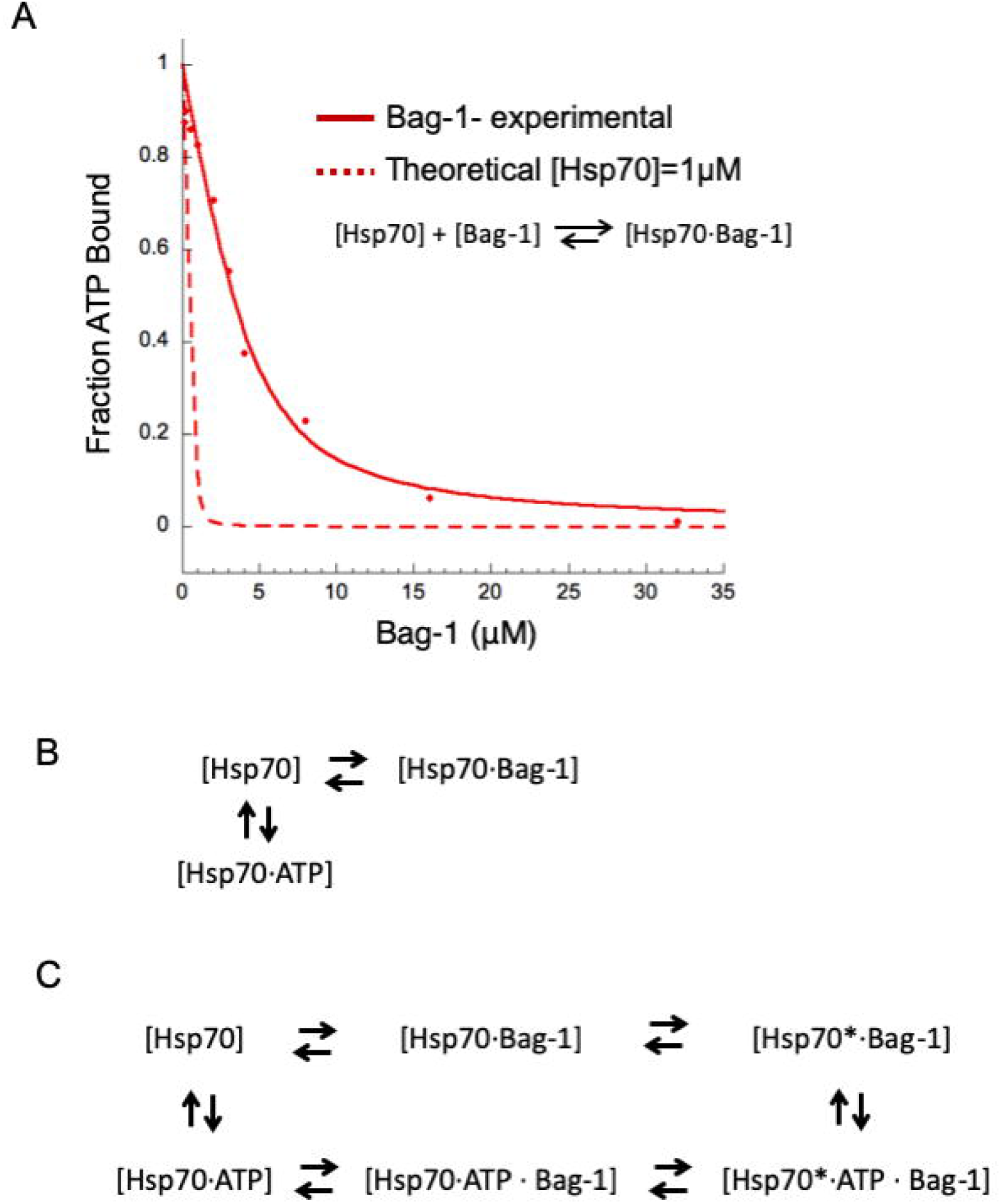
A) Normalized ATP:Hsp70 binding inhibition by Bag-1 as in Fig. S2 but plotted on linear scale and fit with the fraction bound equation (solid line), yielding [A_T_]=4.4µM and K_D_=1µM. Given that [Hsp70_T_] = 1µM, the model for the direct inhibition of ATP:Hsp70 binding by Bag-1 as depicted in (B) does not well describe Bag-1’s mechanism of ATP binding inhibition. Furthermore, the theoretical fit for [Hsp70]=1µM and estimated K_D_=50nM (dotted line) based on the mechanism in (B) over estimates the inhibition potency of Bag-1. (C) Associative displacement mechanism accounting for NEF stabilized conformational state of Hsp70 with low nucleotide affinity (Hsp70*) as proposed for GrpE.

